# Melt Electrowritten Microfiber-Hydrogel Composite Scaffolds for Aligned Muscle Tissue Engineering

**DOI:** 10.1101/2025.08.20.671328

**Authors:** Adam Rauff, Ievgenii Liashenko, Charlotte Lippa, Samuel R. Nightheart, Kimberly A. Jones, Paul D. Dalton, Robert E. Guldberg, Nick J. Willett

**Author notes:** Funding Statement: The authors appreciate funding of the Wu Tsai Human Performance Alliance and the Joe and Clara Tsai Foundation.

## Abstract

Effective regeneration of skeletal muscle with highly aligned fiber architecture remains a significant challenge in tissue engineering. Structural alignment of muscle constructs along with mechanical integrity are crucial for effective engineering of grafts and microphysiological systems. This study engineered a novel composite microfiber-hydrogel platform using melt-electrowriting (MEW) to fabricate high precision microfiber scaffolds from poly(ε-caprolactone). Three MEW scaffold designs (Isotropic, Aligned T with perpendicular reinforcements, and Aligned X with angled fiber bridges) were developed and fabricated into composite scaffolds with collagen hydrogels and seeded with myoblasts. Aligned X scaffolds with cross-bridge reinforcements exhibited enhanced mechanical strength and continuous alignment without structural interruption that led to highly aligned and multinucleated cellular organization. The incorporation of collagen hydrogel in composite constructs improved cell seeding efficiency, viability, and metabolic activity compared to scaffolds alone. All scaffold designs provided fiber reinforcement that prevented hydrogel contraction over extended culture periods. Critically, the Aligned X composite constructs significantly increased myoblast differentiation and myotube maturation, evidenced by increased myosin heavy chain expression and myotube diameter. Overall, this composite microfiber-hydrogel approach provides a scalable, structurally stable, and highly aligned platform tailored for enhanced muscle tissue engineering applications, representing an advancement towards addressing clinical challenges associated with muscle injuries.

## Introduction

Skeletal muscle is essential for locomotion, posture, respiration, and metabolism^1^. Effective regeneration of skeletal muscles following injury is critical for maintaining health and quality of life. Impaired muscle function is classified as a disease and often leads to significant disability, chronic pain, and restricts an individual’s ability to work and participate in recreational activities^2-6^. To address such clinical challenges—primarily arising from aging, injury, or disease—there is a growing interest in regenerative therapies. Tissue engineering of skeletal muscle offers promising solutions, including grafts for volumetric muscle loss (VML) injuries^5,7-9^, and microphysiological systems that enhance our understanding of muscle regeneration and disease^10,11^.

A major barrier to functional muscle regeneration is the disruption of the structural hierarchy that characterizes skeletal muscle tissue^9,12,13^. Skeletal muscle features an organized, multi-scale architecture: at the micron scale sarcomeres form myofibrils; muscle fibers (individual muscle cells) are approximately 100μm in diameter; and fascicles group these fibers into larger functional units (∼30mm)^14^. This hierarchical organization is supported by connective tissue, including the endomysium around fibers, perimysium surrounding fascicles, and epimysium encompassing the entire muscle. The extracellular matrix (ECM) components bear a significant portion of passive loads, with the ECM modulus estimated to be 5-25 times greater than that of muscle cells^15^. Crucially, these structures also facilitate lateral force transmission, enabling efficient mechanical load distribution across parallel aligned structures. The importance of preserving this architecture is underscored by the stark difference in healing outcomes between small and large muscle defects, as regeneration is significantly impaired when tissue loss exceeds a critical threshold^16^. However, current engineered muscle constructs often lack sufficient size, structural alignment, and mechanical integrity to replicate these essential features.

Skeletal muscles experience diverse dynamic and passive loads and possess highly anisotropic mechanical properties, posing significant challenges for engineered constructs^17^. Native muscle tissues are structurally adapted to withstand both active and passive forces, requiring robust mechanical support for stability^14^. Nonetheless, current tissue-engineered muscle constructs fail to fully replicate this complex biomechanical environment. Despite advancements in strategies for cellular alignment, many constructs exhibit poor architectural fidelity and substantial heterogeneity, particularly in thicker tissues^18^. Engineered alignment features are typically 10-to 100-fold larger than the cellular scale, and achieving consistent alignment throughout large-volume constructs remains technically challenging^19^. Moreover, 3D aligned constructs frequently suffer from insufficient mechanical stability, limiting their capacity to effectively transmit force longitudinally along myofibers and laterally through intramuscular connective tissue, both essential for muscle function^20,21^.

Several tissue engineering strategies have attempted to address the need for alignment, but most fall short due to manufacturing limitations^18^. Electrospun fiber membranes are widely used to produce aligned scaffolds with individual fiber diameters typically ranging from 150 nm-2 µm, with varying alignment and mechanical properties^22-26^. However, their high fiber density restricts cell infiltration, typically limiting their application to two-dimensional formats. Additionally, the instability and whipping of the electrospun jet compromises precise control over scaffold architecture. Alternatively, sponge-like scaffolds enable control over porosity - including pore size, orientation, and interconnectivity – to enhance cell infiltration^27^. Yet, increased porosity often reduces mechanical strength, and the achievable alignment remains insufficient for replicating native muscle architecture. Hydrogels delivered via bioprinting or injection methods provide benefits in terms of cell viability and spatial distribution but lack both structural strength and microscale alignment^28-30^. Finally, soft lithography can produce micropatterned films with high fidelity surfaces cues; however, these cues are limited to surface interactions and lack the volumetric architectural control required for engineering thicker tissue constructs^31,32^.

An emerging 3D printing technology called melt electrowriting (MEW) enables the fabrication of microfiber scaffolds with controlled porosity, mechanical integrity, and architectural design over multiple scales. This technology allows the creation of highly anisotropic structures spanning the cellular and tissue scales and supports the manufacturing of large tissue constructs with high porosity^33^. MEW produces scaffolds with fiber diameter typically in the range of 2-50 µm with high precision^34^ while permitting highly intricate architectural designs in large tissue constructs^35,36^. Recently, MEW has demonstrated promise in engineering tendons and ligaments^37^, cardiovascular tissue^38^, and cartilage tissue^39^, yielding constructs that are both highly porous and structurally robust. Additionally, it has been employed to fabricate skeletal muscle scaffolds mimicking the hexagonal fiber morphology characteristic muscle tissue cross-sections^40^. However, the resistance of microfibers to bending forces is poor, as the deformation scales with increases proportionately with the fourth power of the fiber diameter. As a result, scaffolds produced by MEW using microfibers at the cellular scale deform easily, typically necessitating periodic reinforcement with larger fibers/structures^41^. Although reinforcement structures help maintain mechanical stability, their incorporation reduces scaffold porosity and can negatively affect cellular organization. Conversely, removing such reinforcements significantly compromises structural integrity, posing a persistent limitation for tissue-engineered muscle constructs.

In this study, we engineered a composite microfiber-hydrogel platform for skeletal muscle tissue engineering based on MEW and type I collagen hydrogel. Our primary objective was to create large-scale, highly aligned, and mechanically stable muscle constructs capable of promoting muscle cell fusion and differentiation. To address this, we developed a novel microfiber architecture featuring cross-bridge reinforcements, where short diagonal segments connect adjacent parallel fibers that mechanically stabilize the scaffolds. We hypothesized that these composite constructs would support highly aligned cellular organization, enhance myogenic maturation and differentiation, and improve overall structural integrity.

## Materials and Methods

### Microfiber Scaffolds

MEW scaffolds made from 10 µm microfibers were 3D-printed using of poly(ε-caprolactone) (PCL) (PURAC PC12, Corbion Inc.). Fibers were deposited into a total of 20 layers with three different designs while keeping the total amount of extruded material consistent. The three scaffold architectures include 1) “Isotropic” 5-fiber morphology (Figure 1A,D,G), 2) “Aligned T” fibers with perpendicular running reinforcements (Figure 1B,E,H), and 3) Aligned X fibers with cross-bridge reinforcements at an angle between 30-60^°^ and semi-randomly dispersed throughout the scaffold plane (Figure 1C,F,I). The scaffolds were fabricated in a circular shape with an overall diameter of 13.0 mm. The isotropic scaffold was previously used for an organoid model of skin^42^, and contains spacing between each parallel track of fibers of 150 *μ*m, with fibers running at 0°, 36°, 72°, 108°, and 144°(Figure 1A,D,G). The aligned T scaffold architecture was previously used for engineered articular cartilage^39^, and employed 100 µm between aligned fiber walls and 1.5 mm between reinforcing perpendicular walls (Figure 1B,E). The aligned X microfiber architecture had 100 µm spacing between aligned fiber walls while cross-bridge reinforcements were accomplished by offsetting fiber directions by ±3° from the principal alignment axis in 8 of the 20 layers comprising each scaffold. The offset caused deposition of a fiber on previously laid material and led to “fiber bridging”^43^ to the adjacent fiber wall as the planar distance of the extruded jet and the original fiber wall increased. The number of layers was chosen by testing 3 different configurations with 5, 8 or 10 offset layers. The Aligned X scaffold with 8 bridging fiber layers interlaced uniformly with the aligned layers and qualitatively yielded the best combination of maintaining shape and porosity (Figure 1C, Supplementary Figure 3). All scaffolds included two reinforcement rings each from 4 layers of large, 70 µm diameter, fibers along their perimeter which improved construct handling under wet and dry conditions by resisting folding, bending, and kinking (Supplementary Figure 1). Scanning Electron Microscopy (SEM) was used to visualize and qualitatively compare the three scaffold designs. Scaffolds were sputter-coated with 7 nm of titanium prior to imaging with an SEM Everhart–Thornley detector (ThermoFisher Apreo 2, USA).

**Figure 1.**
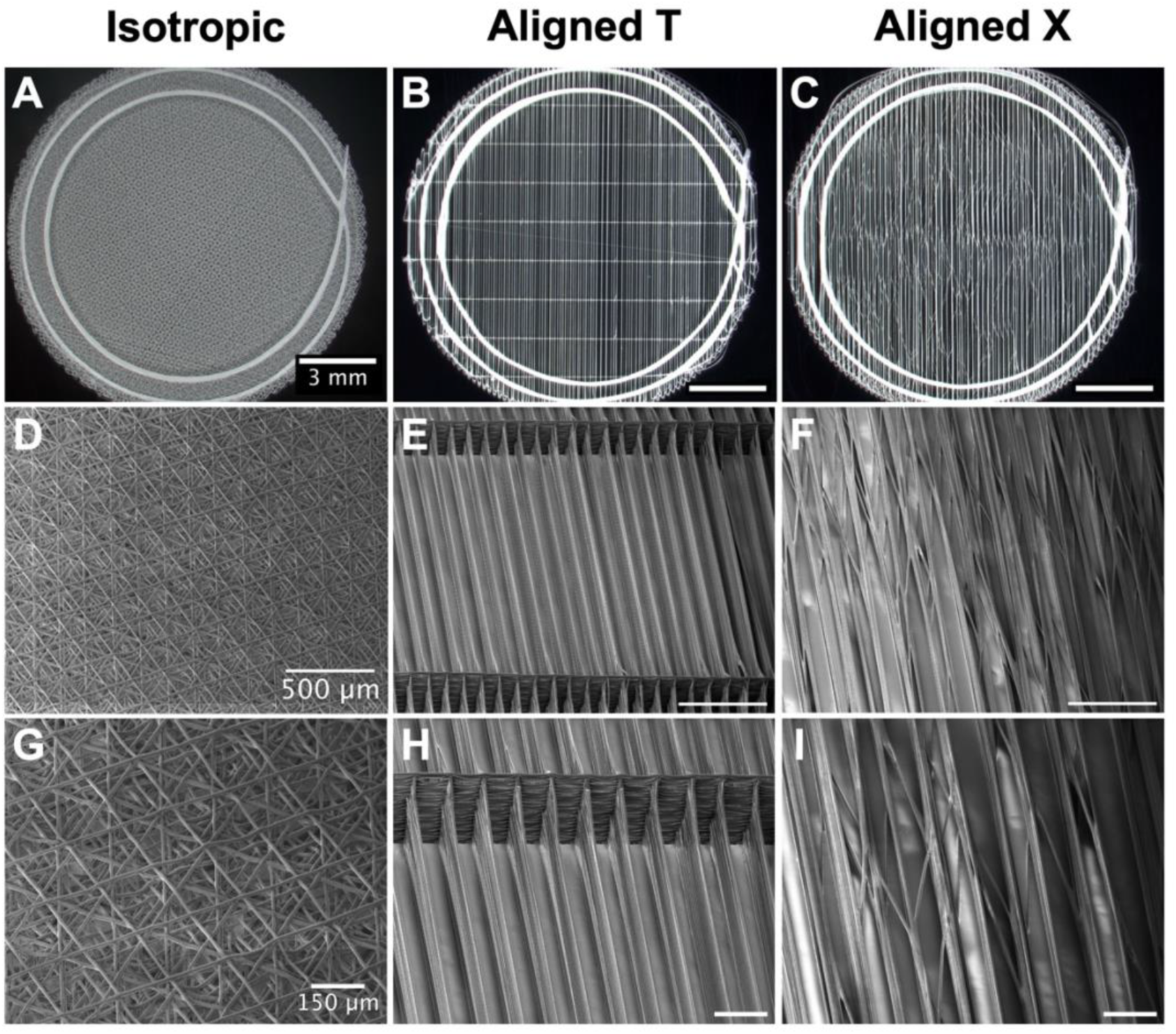
Three microfiber scaffold designs were manufactured using melt-electrowriting (MEW) - a 3D printing technique. The scaffold architectures were imaged with a light microscope comprising **A)** Isotropic 5-fiber family, **B)** Aligned T fibers with perpendicular reinforcements, and **C)** Aligned X fibers with angled cross-bridge reinforcements. All scaffolds had a circular shape with an overall diameter of 13.0 mm that fit in a 24 wellplate. Details of the scaffold architecture were visualized from scanning electron microscopy (SEM). **D)** The Isotropic scaffold contained spacing between each parallel track of fibers of 150 μm. with fibers running at 0°. 36°. 72°, 108°, and 144°. **E)** The aligned T scaffold architecture employed 100 pm between aligned fiber walls and 1500 pm between reinforcing perpendicular walls. **F)** The aligned X microfiber architecture was designed with 100 pm between aligned fiber walls and 8 bridging fiber layers interlaced uniformly with the aligned layers. **G-l)** Higher magnification SEM images show the precision of manufacturing features on a small scale.

Scaffolds were fabricated using a custom-built MEW printer using the following processing and instrument parameters: a melt temperature of 75°C was used to extrude to an electrically grounded 22 G nozzle (with 6.3 mm length) protruding 1 mm from the printhead. Nozzle-to-collector distance was fixed to 4 mm, and scaffolds were printed onto 1 mm thick glass microscopy slides resting on a metal collector with an applied voltage potential of 5.5 kV. Fibers with 10.0 µm were achieved by applying 0.17 bar pressure and 500 mm/min collector translation speed. Reinforcement rings along scaffold perimeter with 70.0 µm fiber diameter were printed by applying 3.0 bar pressure and 65 mm/min collector translation speed. Microfiber scaffolds with reinforcement rings were released from the glass slides by wetting with ethanol solution.

### Mechanical Testing

Tensile properties of dry, acellular microfiber scaffolds were evaluated using a TA Electroforce 3220 mechanical tester equipped with a 1000g load cell. Rectangular samples were carefully cut from circular scaffolds using surgical scissors to a target width of 4.0 mm. Actual measured widths of the scaffolds were not significantly different across the groups, averaging 4.63±0.55 mm, 4.25±0.94 mm, and 4.30±0.51 mm for the Isotropic, Aligned T, and Aligned X groups respectively. Scaffold lengths were measured between clamps upon mounting and were not significantly different with mean lengths of 9.47±0.90 mm, 8.25±0.99 mm, and 8.50±0.84 mm. Tests were performed to failure with a strain rate of 0.2 mm/s with the recorded signal of force data recorded at a minimum frequency of 10Hz. Post-testing analysis utilized custom Matlab (MathWorks) code to calculate Young’s modulus, yield stress, and yield strain. Nominal stress was computed by dividing the measured force by the sample’s cross-sectional area, assuming uniform thickness due to consistent fiber layering and total scaffold mass. Strain was computed relative to initial sample length. Stress-strain curves were smoothed with a three-position moving window filter. Elastic modulus and yield parameters were numerically identified using the derivative of the stress-strain curves: the elastic region was defined as a drop of 20% of the peak slope, while yield points corresponded to the location slope decreased to 20% of the peak slope following linear elasticity. Young’s modulus was determined by linear regression within the elastic region, and the yield stress/strain were recorded at the yield point.

### Seeding Muscle Constructs with Microfiber Scaffolds

Composite muscle constructs were fabricated by embedding microfiber scaffolds within collagen hydrogels containing suspended cells. Microfiber scaffolds were sterilized in 70% ethanol for a minimum of 24 h, rinsed three times with 1X phosphate-buffered saline (PBS), and incubated in growth medium for a minimum of 12 hours at 37°C. C2C12 myoblasts cells (ATCC, cat. CRL1772) were suspended in type I collagen solution (Advanced Biomatrix; 1.5 mg/mL), at a density of 400,000 cells/mL, unless otherwise specified. 100μL of collagen-cell suspension was pipetted onto scaffolds placed within ultra-low attachment 24 well plates (Costar, Fischer Scientific) and polymerized at 37°C.

Scaffold-only constructs, used as controls, were identically prepared and seeded with an equivalent total cell number (40,000 cells/scaffold). C2C12 myoblasts were directly pipetted onto microfiber scaffolds rather than within a collagen gel. For metabolic activity and DNA quantification assays, a higher seeding density of 800,000 cells/mL was additionally tested, with corresponding scaffold-only constructs seeded at 80,000 cells/scaffold for this condition.

### Cell Culturing Conditions

Constructs were cultured initially in growth medium comprised of Dulbecco’s Modified Eagle Medium (DMEM), 10% Fetal Bovine Serum (FBS), and 1% penicillin and streptomycin for 5 days. The medium was switched to differentiation medium (DMEM, 2% horse serum, and 1% penicillin/streptomycin) for 3 additional days, with media changes on odd days. For differentiation and maturation assays, the total culture duration was extended to 16 days.

### Quantification of Cellular Alignment

Cellular alignment in composite microfiber-hydrogel constructs (Isotropic, Aligned T, Aligned X) was evaluated after 8 days of culture. Additional control groups included monolayer culture on tissue culture plastic (5,000 cells/cm^2^), hydrogel-only (collagen hydrogel without a scaffold), and electrospun mesh (ES Mesh) without a hydrogel. Due to rapid confluency and contraction, monolayer and hydrogel only controls were fixed after 4 days. Hydrogel-only controls were seeded with 300μL of collagen-cell solution and identical cell concentration to composite constructs to ensure a continuous layer covering the entire area of the well in the absence of the scaffold.

Images were acquired with LAS X software using a Leica Thunder (Leica Microsystems) inverted microscope with computational clearing to reduce background noise. Volumetric Z-stack images were acquired at three randomized nonoverlapping locations using the 5X objective (n=3 technical replicates). The DAPI channel was imaged using the 390nm LED at 40% illumination with a 460nm emission filter, while the F-actin channel was imaged using the 510nm LED at 50% illumination with a 535nm emission filter. All images were acquired with 100ms exposure time. The maximum intensity projections of the Z-stack were used for analysis. Images of the F-actin channel were pre-processed by converting it to an 8-bit image, enhancing the contrast using histogram equalization, and applying a median filter with a radius of 3.0 pixels. Then a directional analysis based on the Fourier transform was applied to quantify a directional histogram encompassing 180 degrees with 180 discrete bins. The Gaussian distribution contains a single peak, and its goodness-of-fit was used as a uniaxial alignment index. The regional variance of alignment was quantified using the standard deviation of the mean direction of each technical replicate. Images were analyzed and prepared for publication using ImageJ^44^.

### Immunostaining and Fluorescence Microscopy

Constructs were fixed with 4% paraformaldehyde (Fisher Scientific) for 30 minutes, permeabilized in 0.1% Triton-X-100 (Millipore Sigma) solution for 20 minutes, blocked in 2% bovine serum albumin (Millipore sigma) for 40 minutes, with phosphate buffered solution (PBS) rinses between each step. Analysis of cellular alignment utilized DAPI (Biolegend) and AlexaFluor 488 phalloidin (ThermoFisher) to stain nuclei and cellular cytoskeleton. The stains were diluted to 300 nM for DAPI and 400X for phalloidin in blocking solution and incubated for 1 h at room temperature. Analysis of differentiation and maturation utilized a primary antibody to stain myosin heavy chain (MHC) (Abcam, ab91506) with 1:500 dilution incubated in 4C° overnight. Secondary staining included DAPI, AlexaFluor 568 phalloidin to stain nuclei and cellular cytoskeleton, and 1:200 dilution of AlexaFluor 488 IgG as a secondary antibody (Abcam, ab150081). Samples were incubated in secondary antibodies diluted in blocking solution for 60 minutes at room temperature.

### Hydrogel Contraction

The contraction of samples was measured by quantifying the retained scaffold area and comparing it with hydrogel-only controls at days 4 or 8. Constructs were washed with 1X PBS and stained (0.01% toluidine blue) at 0.5 mL/sample for 5 minutes, and then washed again with 1X PBS. A stationary digital camera (Canon) with an ultrashort focal length lens was used to acquire images, which were processed and analyzed in ImageJ ^44^. The area was calibrated against standard 24-well dimensions (15.6 mm) and measured for retention percentage compared to the initial area.

### Cellular Viability

Cellular viability was assessed 24 h post-seeding using propidium Iodide (PI, ThermoFisher) for dead cells and DAPI for total nuclear count. Z-stack images were acquired at three randomized nonoverlapping locations using the 10X objective (n=3 technical replicates). Constructs with sampled images with an average of less than 30 nuclei per image were excluded from the analysis to avoid inaccurate representation of viability. The images were pre-processed using a median filtered, and the PI channel was thresholded using the positive-control baseline to remove background noise. DAPI images were thresholded manually by visual inspection. The images of each sample were then processed together to compute the Manders’ Coefficient (M2), to measure the degree of overlap between the two channels^45^. Viability was defined by V=(1-M2)*100. This ensured the PI signal was specific to a DAPI-positive nuclei. Image data were analyzed using ImageJ^44^ and the Coloc2 algorithm.

### Cellular Proliferation

Proliferation was measured using the Click-iT® EdU cell proliferation assay (Thermofisher). Constructs were incubated with 5-ethynyl-2′-deoxyuridine (EdU) at 10 μM in culture medium at Day 0, fixed after 24 h, stained according to manufacturer guidelines. Briefly, fixed constructs were permeabilized in 0.1% Triton X-100 solution for 20 minutes, then counterstained using anti-EdU reaction solution for 30 minutes at room temperature. Nuclei were counterstained with DAPI, and images were acquired using a Nikon spinning disk confocal microscope with a 10X objective. Volumetric Z-stack images were acquired at three randomized non-overlapping locations, with exclusion criteria of less than 30 total nuclei. Proliferation percentages were quantified based on EdU/DAPI colocalization.

### Metabolic Activity and DNA Quantification

Construct metabolic activity (alamarBlue^™^ assay; Thermofisher, A50100) and total DNA content (picoGreen assay; Thermofisher, P7589) were assessed 24 h post-seeding. Two cell seeding concentrations were studied, including 400,000 cells/mL and 800,000 cell/mL, comparing composite and scaffold-only constructs while keeping the total cells consistent across the constructs without hydrogel. Isotropic scaffold design represented the microfiber scaffolds, and with control groups consisted of hydrogel-only without scaffold (Gel-only) and monolayers cultured on standard tissue culture plastic. After 24 h of culture, cells were washed with PBS and incubated in 10% alamarBlue for 2 h. The media was analyzed with a microplate reader with technical duplicates and then averaged. The scaffolds were then digested in 0.5 mg/mL papain (Millipore Sigma, 10108014001) at 65°C overnight. After 16 hours of incubation, samples were stained with PicoGreen solution, analyzed with a microplate reader, and calibrated against standard dilution curve.

### Myoblast differentiation and maturation

Differentiation was evaluated by quantifying myosin heavy chain (MyHC) expression after 16 days, using immunohistochemical staining. Composite constructs were seeded at a concentration of 800,000 cells/mL, changed to differentiation medium on Day 5, and fixed on day 16. Samples were imaged on Zeiss LSM 880 confocal microscope with a 20X objective. The acquisition settings included laser power of 3.0%, 2.4%, and 2.0% for the for the blue, green, and red excitation lasers, a gain of 849,578, 617.208, and 759.740 for the blue, green, and red emission channels, and a pixel dwell time of 2.576•10^-7^ seconds. Images comprised three z stacks per sample at random non-overlapping locations within the constructs, encompassing a depth of the entire sample. The stacks were median filtered using a 3-element cubed structuring element, and maximum intensity projections were obtained. Outcome measures included relative area fraction, myotube count, myotube diameter, and mean fluorescence intensity (MFI). The MFI was acquired from the raw images of the MyHC channel by computing the mean intensity of all pixels. The analysis for the other outcomes followed established protocols from previous publications^46^. The green channel representing MHC expression was processed using localized histogram equalization (CLAHE). The images were then manually thresholded by choosing the triangle criteria as an initial guess and increasing the threshold until the background scatter was fully removed. Then, images were converted to a mask, underwent the morphological imaging operation of opening to remove small objects and filling holes. Myotubes were identified using the Analyze Particles analysis with a lower cutoff of 400 μm. This enabled measurement of the relative area fraction inhabited by myotubes and the number of myotubes within the imagining region. The myotube diameter was then acquired using 5 measurements of the segmented myotubes with no more than 1 measurement per myotube.

### Statistical Analyses

The data were analyzed by a one-way ANOVA and a Tukey’s post-hoc analysis (95% CI). A two-way ANOVA was used in the hydrogel contraction, nuclear count, cellular viability, proliferation, metabolic activity, and DNA content, to analyze the effects of two independent variables along with Sidak’s multiple comparisons tests. Statistical significance was determined at p=0.05, and 0.10>p>0.05 were displayed on graphs. All statistical analyses were conducted using GraphPad Prism version 10.0 (GraphPad Software, MA).

## Results

### MEW Enables Fabrication of Large, Aligned, and Stable Scaffolds with Micron-scale Fibers with Unique Architecture

MEW enabled the fabrication of large scaffolds with 13.0 mm diameter, using high precision 10 µm filament fibers (Figure 1). Each scaffold incorporated two reinforcement rings, fabricated from larger fibers (∼70 µm diameter) along their perimeter, which substantially enhanced handling under wet and dry conditions by resisting folding, bending, and kinking (Figure 1A-C, Supplementary Figure 1). SEM confirmed MEW’s capacity to manufacture large tissue constructs with micron-sized structural precision (Figure 1D-I). The three scaffold architectures – Isotropic, Aligned T, and Aligned X – demonstrated the versatility of the MEW technology for tailoring the scaffold microenvironments in engineered tissues. The Aligned X architecture featured intermittent diagonal reinforcements (angles of 30-60^°^), providing superior stability compared to perpendicular reinforcements. Specifically, aligned fibers of the Aligned T scaffold architecture (with perpendicular reinforcements) exhibited deformation between the supports, forming a wavy pattern, while aligned fibers in Aligned X scaffolds maintained their integrity and alignment (Supplementary Figure 2, Supplementary Videos 1-3). Scaffolds with aligned fibers but no reinforcements failed to maintain their shape in aqueous conditions (Supplementary Figure 1C). By contrast, the aligned fibers of aligned T were interrupted by perpendicular reinforcements (Figure 1E,H). The crossbridge reinforcement enabled highly aligned design features over 10 mm of length without a physical interruption (Figure 1C).

### Microfiber Scaffold Architecture Modulates Bulk Mechanical Properties

The mechanical properties of microfiber scaffolds were driven by fiber architecture. Tensile testing along the preferred fiber orientation revealed significant differences in the young’s modulus, yield stress, and yield strain across the scaffold designs (Figure 2). The Young’s modulus had two-fold difference from the lowest to the highest modulus groups, from 3.89 ±0.575 MPa in the Isotropic group, 6.35 ±0.800 MPa in the Aligned T, and 10.1 ±1.62 MPa in the Aligned X group (Figure 2A), with statistically significant differences between groups (p<0.004). Yield stress of Aligned X and Aligned T scaffolds required significantly more load before plastic deformation, compared with Isotropic scaffolds (p<1·10^-6^). Conversely, yield strain indicated the greatest elastic deformation in the Aligned T scaffolds and the lowest in Aligned X (Figure 2C), with significant differences among all groups (p<0.0350). These results underscore the scaffold architecture’s decisive role in modulating bulk mechanical properties. Additionally, scaffold stiffness decreased after incubation with media over 11 days, irrespective cellular presence (Supplementary Figure 4), following a similar observation for MEW scaffolds used as a skin model^42^.

**Figure 2.**
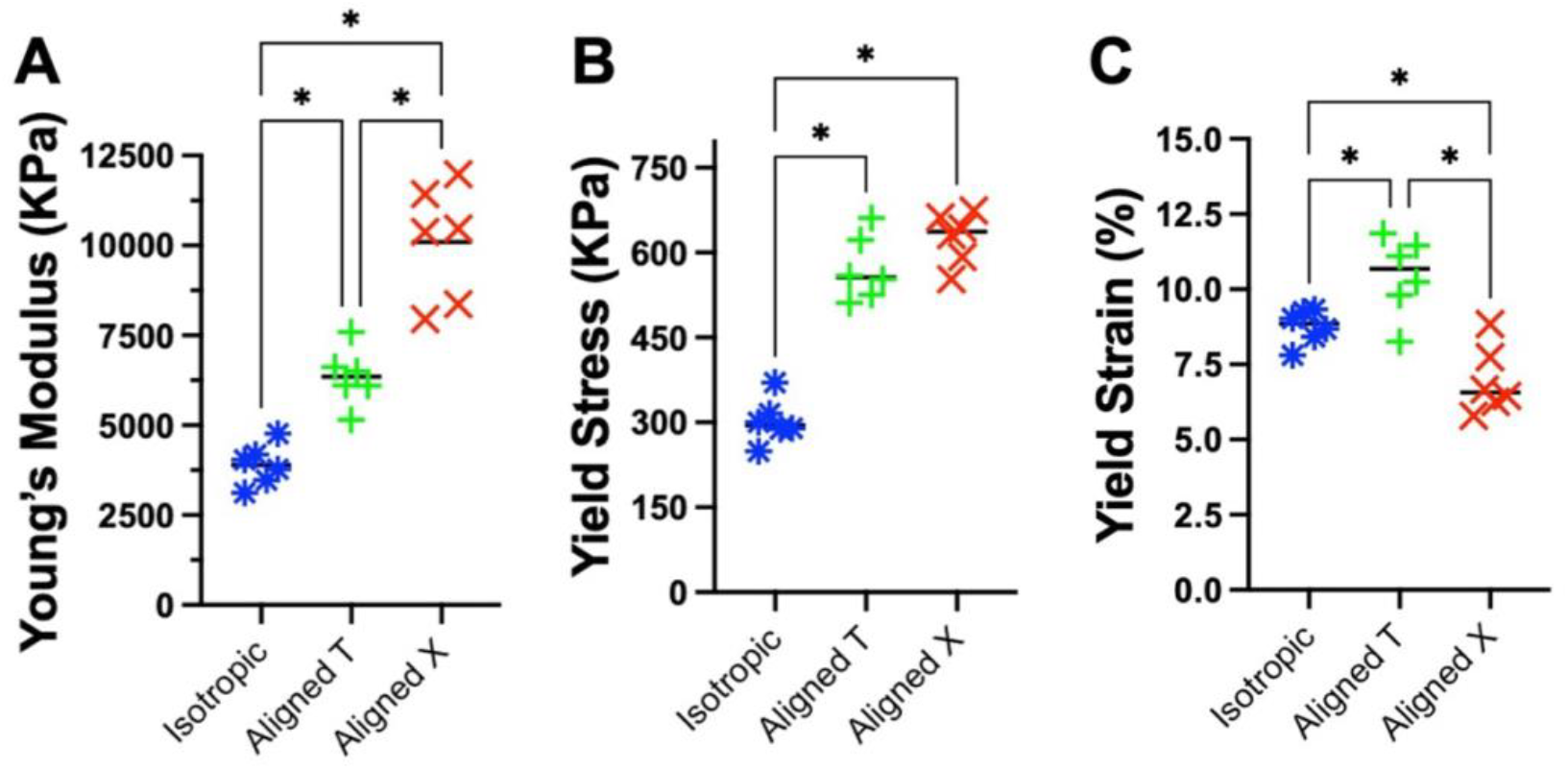
Tensile strength of the dry, acellularized, microfiber scaffolds was assessed comparing the three architectural designs – Isotropic, Aligned T, and Aligned X. Tensile testing along the preferred fiber orientation revealed significant differences scaffold designs. **A)** Graph comparing the Young’s modulus of the three scaffold designs shows the Aligned X had the highest modulus that was more than double that of the Isotropic group. All three groups were significantly different from each other. **B)** Graph examining the yield stress across the scaffold designs showed the Isotropic group required significantly less load to reach plastic deformation. **C)** Graph depicting yield strain showed significant differences across all groups with the Aligned T enabling the most deformation prior to plastic deformation. The solid black line represents the mean of each distribution. Statistical significance was determined at p<0.05 with one asterisk. A one-way ANOVA was used with a Tukey’s post-hoc.

### Aligned Scaffold Architecture Induced Cellular Alignment

Composite microfiber-hydrogel constructs significantly influenced cellular alignment. By day 8, myoblasts co-localized and aligned closely with the microfiber scaffolds, adopting the scaffold’s geometry at both the cellular and tissue scales (Figure 3A-C, G-I; Supplementary Figure 5). Constructs with Aligned T and Aligned X scaffolds displayed substantially higher aligned cellular organization over the isotropic design. The aligned T group had distinct cellular structures along the perpendicular reinforcing fibers (Figure 3E,H, white arrows). The perpendicular reinforcements (Figure 1H) impeded formation of continuous cellular bodies across the wall (Figure 3H). The uniaxial alignment index of Aligned T and Aligned X constructs was significantly higher than the Isotropic (p<1·10^-6^) and all control groups (p<0.0030, Figure 3J). The gel-only group had higher alignment index than the Isotropic (p=0.0069) and electrospun constructs (p=0.0092), potentially resulting from the axis of folding due to hydrogel contraction at Day 4 (Supplementary figure 6C). The heterogeneity of cellular organization was measured using the spatial variance of the mean direction. The mean variance of the Aligned X and Aligned T groups were 0.848^°^ ±0.509^°^ and 2.908^°^ ±3.036^°^ respectively and was significantly lower than 2D monolayer cultures (p<0.0304) and electrospun groups (p<0.0086, Figure 3K).

**Figure 3.**
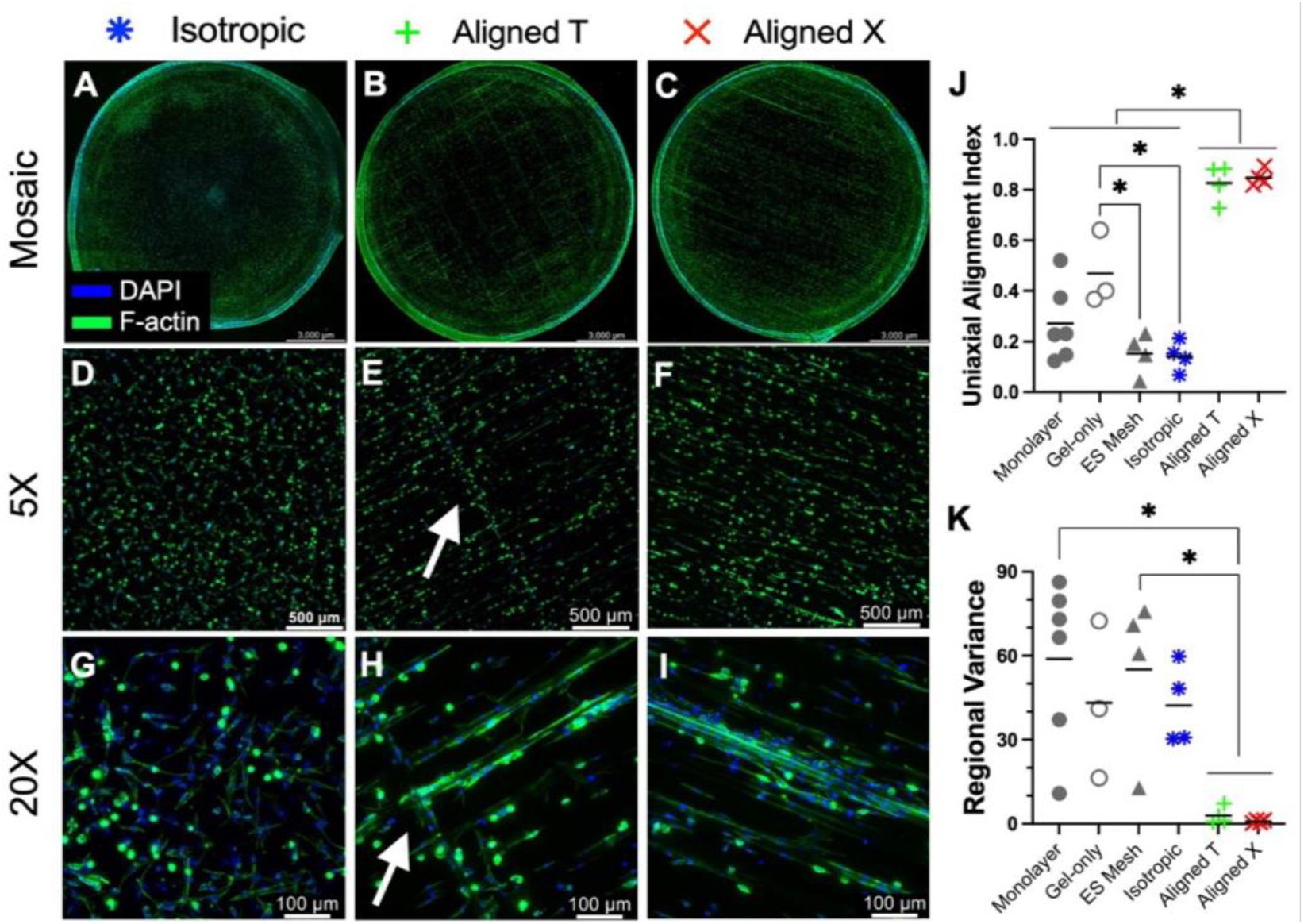
Aligned microfiber architecture induced cellular alignment in composite constructs. **A-C)** composite muscle constructs cultured to Day 8 and visualized via immunofluorescence microscopy showing Isotropic (A), Aligned T (B), and Aligned X (C) scaffold designs. Images were acquired with a 5X objective and stitched together to show the entire scaffolds (scale bar - 3.0mm). **D-F)** Individual 5X images demonstrating cellular organization on each scaffold design (scale bar – 500 µm). The Aligned T design had distinct cellular structures along the perpendicular reinforcements (E, white arrow). **G-I)** Higher magnification images acquired with a 20X objective (scale bar – 100 µm). The Aligned T design had distinct cellular structures along the perpendicular reinforcements, and appeared to impede continuous cellular structures (H, white arrow). The DAPI channel was imaged using the 390nm LED at 40% illumination with a 460nm emission filter, while the F-actin channel was imaged using the 510nm LED at 50% illumination with a 535nm emission filter. All images were acquired with 100ms exposure time. **J)** Comparison of the uniaxial alignment index across composite constructs and three control groups including monolayers, hydrogel-only (Gel-only), and electrospun mesh (ES Mesh) without hydrogel. Composite constructs with the Aligned T and Aligned X scaffold designs had significantly higher alignment than the Isotropic constructs and the three control groups. **K)** Comparison of the regional variance of mean direction amongst three technical replicates from different location in each sample. The Aligned T and Aligned X constructs had the lowest spatial variance and were significantly lower than the monolayer and ES mesh groups. This demonstrates more spatial homogeneity in the aligned scaffold architectures. The solid black line represents the mean of each distribution. Statistical significance was determined at p<0.05 with one asterisk. A one-way ANOVA was used with a Tukey’s post-hoc.

### Scaffold Reinforcement Prevents Cell-mediated Collagen Hydrogel Contraction

Microfiber scaffolds effectively counteracted cell-mediated hydrogel contraction. Collagen hydrogels without reinforcement (Figure 4A) contracted substantially more by day 4 than composite constructs with Isotropic (Figure 4B), Aligned T (Figure 4C), or Aligned X (Figure 4D) scaffold designs. By day 8, hydrogel-only constructs further contracted and assumed an irregular shape (Figure 4E-H). The amount of hydrogel contraction was attenuated in MEW fiber-reinforced hydrogels, maintaining an average of 87.0%, 84.6%, and 83.64% of area retention on day 4, and 82.7%, 82.7%, and 81.9% of area retention on day 8 for Isotropic, Aligned T, and Aligned X constructs respectively (Figure 4I). The contraction of the hydrogel-only constructs differed significantly from day 4 (33.1%) to day 8 (13.2%) with p<0.000001. By contrast, the composite groups mostly maintained their area from day 4 to day 8, with only the Isotropic group statistically decreased in hydrogel area retention from day 4 (87.0%) to day 8 (82.67%) with p=0.0093. The Aligned T group had internal fissures on day 8, with the hydrogel phase separating along the perpendicular reinforcements (Figure 4G). All constructs had darker toluidine blue staining on day 8, suggesting compositional change in the material (Figure 4F-H). These data demonstrate the microfiber scaffolds reinforced the structural stability of composite engineered muscle constructs by resisting cell-mediated contraction of collagen hydrogel.

**Figure 4.**
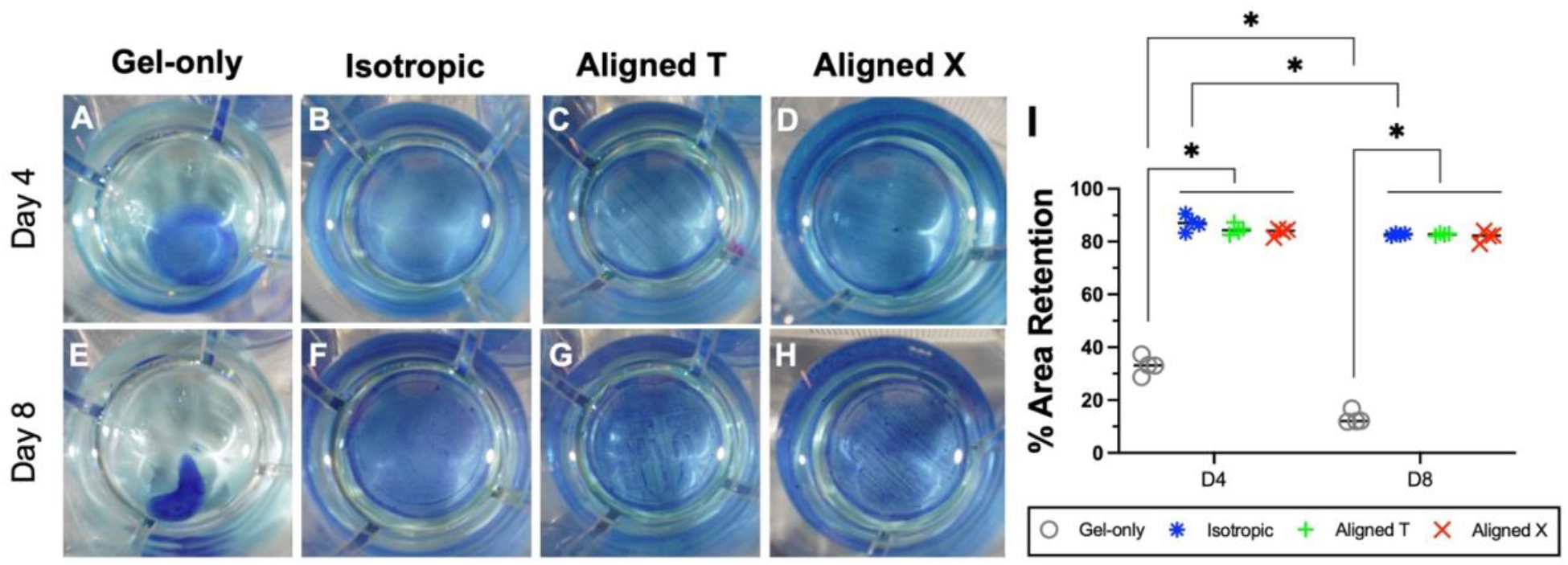
The microfiber scaffolds provide a fiber reinforcement to composite constructs that prevented cell-mediated hydrogel contraction. **A-D)** Representative images of myoblasts cultured for 4 days on hydrogel-only (A), and composite microfiber-hydrogel constructs with Isotropic (B), Aligned T (C), and Aligned X (D) scaffold designs. Images were acquired using a DSLR camera with the constructs in 24 wellplates. The diameter of each well in a 24 wellplate is 15.6mm. **E-H)** Representative images of constructs cultured for 8 days with groups consistent column-wise with (A-D). **I)** Graph portraying the area retention of each group across the two timepoints. The hydrogel-only group (Gel-only in gray circles) had significantly less hydrogel area retained compared with composite constructs. The hydrogel-only group also had lower area retention on day 8 compared with day 4. The three composite constructs groups were not different from each other on day 4 or day 8. The Isotropic group had statistically decreased in hydrogel area retention from day 4 (87.0%) to day 8 (82.67%). The solid black line represents the mean of each distribution. Statistical significance was determined at p<0.05 with one asterisk. A two-way ANOVA was used with Sidak’s multiple comparisons test.

### Incorporation of a hydrogel enhanced metabolic activity and improved cellular seeding efficiency in muscle constructs

The inclusion of hydrogel within composite constructs improved metabolic activity and cell-seeding efficiency. Scaffold-only constructs showed no difference in metabolic activity across the two seeding concentrations (p=0.7708), while composite constructs had increased metabolic activity with higher cell seeding concentration (p=0.0139, Figure 5A). Analysis of DNA mass revealed consistent results. Composite constructs had increased cell DNA (p=0.0303) with increasing cell seeding density, while the scaffold-only group did not show a difference (p=0.6216, Figure 5B). Furthermore, a hydrogel-only group with the same levels of cell seeding concentrations revealed increasing metabolic activity and DNA mass with increasing cell density (Supplementary Figure 6). The high-density composite samples had increased DNA compared with the scaffold-only low density (p=0.0410). These results indicate that incorporation of the hydrogel significantly enhanced the number of cells and metabolic activity.

**Figure 5.**
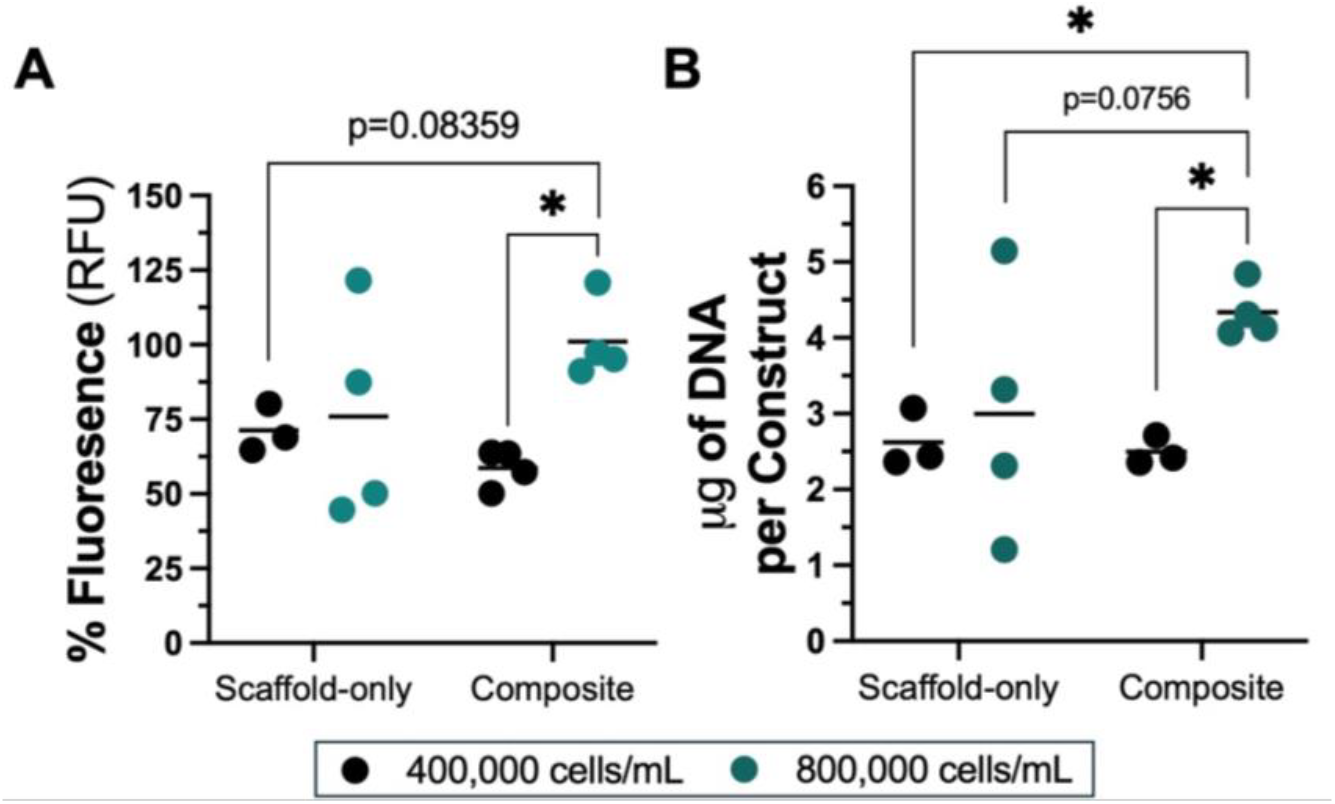
Incorporation of collagen hydrogel enhanced metabolic activity and improved cell seeding efficiency. **A)** Graph showing the results of alamarBlue assay comparing scaffold-only constructs and composite constructs 24 hours post seeding. The groups had two different effective seeding densities of 400,000 and 800,000 cells/mL. The composite constructs had increased signal from the lower to the higher seeding density, while the scaffold-only groups were not different across the seeding densities. **B)** Graph showing the results of picoGreen assay of the same groups in (A). Composite constructs had increased DNA mass with increased cell density, whereas the scaffold-only groups were not different from each other. The solid black line represents the mean of each distribution. Statistical significance was determined at p<0.05 with one asterisk. A two-way ANOVA was used with Sidak’s multiple comparisons test.

### Incorporation of collagen hydrogel improved cell retention and viability of composite constructs across all scaffold designs

The amount of cells retained in muscle constructs and their viability were examined across scaffold-only and composite constructs using nuclear counts and cellular viability. The collagen hydrogel enabled increased number of cells seeded onto constructs with increased viability (Figure 6). A two-way ANOVA showed composite constructs had significantly higher number of nuclei compared with scaffold-only samples (p<0.000001), while the effect of scaffold design was not significant (p=0.3876, Figure 6A). Scaffold-only constructs of Aligned T and X designs had significantly less nuclei than all scaffold design groups of composite constructs (Figure 6A). Composite constructs were consistent with a hydrogel-only group (data not shown), and there were no significant differences between any scaffold designs, suggesting the difference nuclear count is due to the presence of hydrogel (Figure 6A). Moreover, cellular viability was significantly increased in composite constructs compared with scaffold-only constructs (p=0.0060), while the scaffold design architecture did not influence viability (p=0.6154, Figure 6B). The viability of composite constructs was consistent with Hydrogel-only controls (data not shown), suggesting the hydrogel enabled increased cellular viability. Still, cells colocalized with and conformed to the scaffold geometry in composite constructs, suggesting the microfiber scaffolds did affect cellular behavior (Supplementary Figure 5). Furthermore, the proliferation of cells, measured by EdU incorporation, was increased in composite constructs compared with scaffold-only constructs (p=0.0230), and scaffold design also influenced proliferation (p=0.0439, Figure 6C).

**Figure 6.**
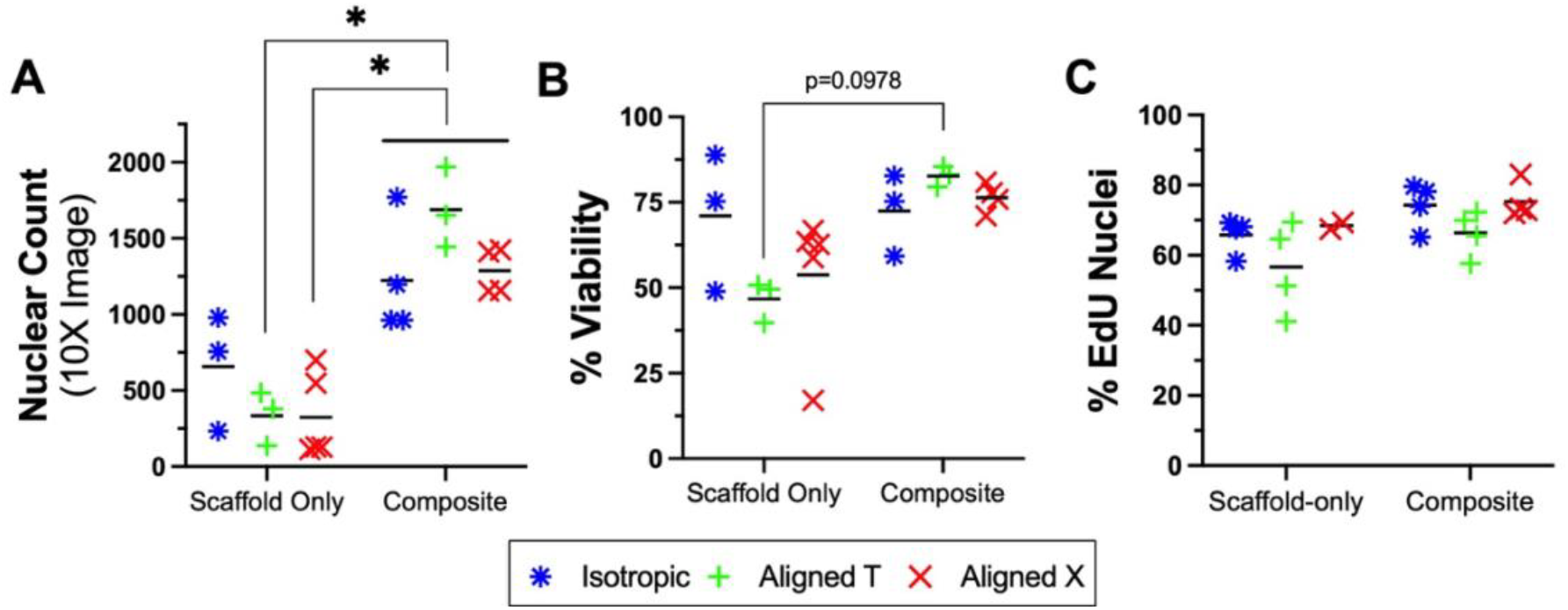
Incorporation of collagen hydrogel enhanced cell retention and increased viability in composite constructs. The analyses compared composite and scaffold-only constructs 24 hours post seeding across Isotropic, Aligned T, and Aligned X scaffold designs. **A)** Graph showing the nuclear count. A two-way ANOVA revealed composite constructs had significantly higher number of nuclei compared with scaffold-only samples (p<0.000001), while the effect of scaffold design was not significant (p=0.3876). **B)** Analysis of cellular viability revealed significant difference across composite and scaffold-only constructs (p=0.0060), while the scaffold design did not influence viability (p=0.6154). **C)** Analysis of cellular proliferation revealed increased proliferation in composite constructs compared with scaffold-only constructs (p=0.0230), and that scaffold design influenced proliferation (p=0.0429). The solid black line represents the mean of each distribution. Statistical significance was determined at p<0.05 with one asterisk. A two-way ANOVA was used with Sidak’s multiple comparisons test.

### Composite Constructs with Aligned X Microfiber Scaffold Increased Myotube Formation

Myotube formation and maturation significantly improved in composite constructs with the Aligned X scaffold. Expression of myosin heavy chain (MyHC) protein was compared across Isotropic, Aligned T, and Aligned X composite constructs seeded with C2C12 myoblasts (Figure 7). The Aligned X constructs exhibited larger diameter multinucleated myotubes compared with the Isotropic and Aligned T groups (Figure 7A-F). The area fraction of myotubes in Aligned X constructs were significantly increased from Isotropic (p=0.0151) and nearly significant compared with Aligned T constructs (p=0.0581, Figure 7G). Together, the data suggest increased expression of MyHC and increased cell differentiation in composite constructs with Aligned X microfiber design. Moreover, the Aligned X constructs also displayed increased maturation. The myotube diameter was significantly higher in the Aligned X group compared with Isotropic (p=0.0007) and Aligned T (p=0.0098) (Figure 7I). The number of myotubes in Aligned T and X constructs significantly increased compared with the Isotropic group (p=0.0003, and p=0.0007 respectively), while there was no difference between Aligned X and Aligned T (p=0.5157, Figure 7J). Moreover, the Aligned T myotubes did not cross over perpendicular reinforcements (white arrow, Figure 7E). Similarly, the Isotropic group had sparse presence of multinucleated structures that were interrupted by reinforcing microfibers.

**Figure 7.**
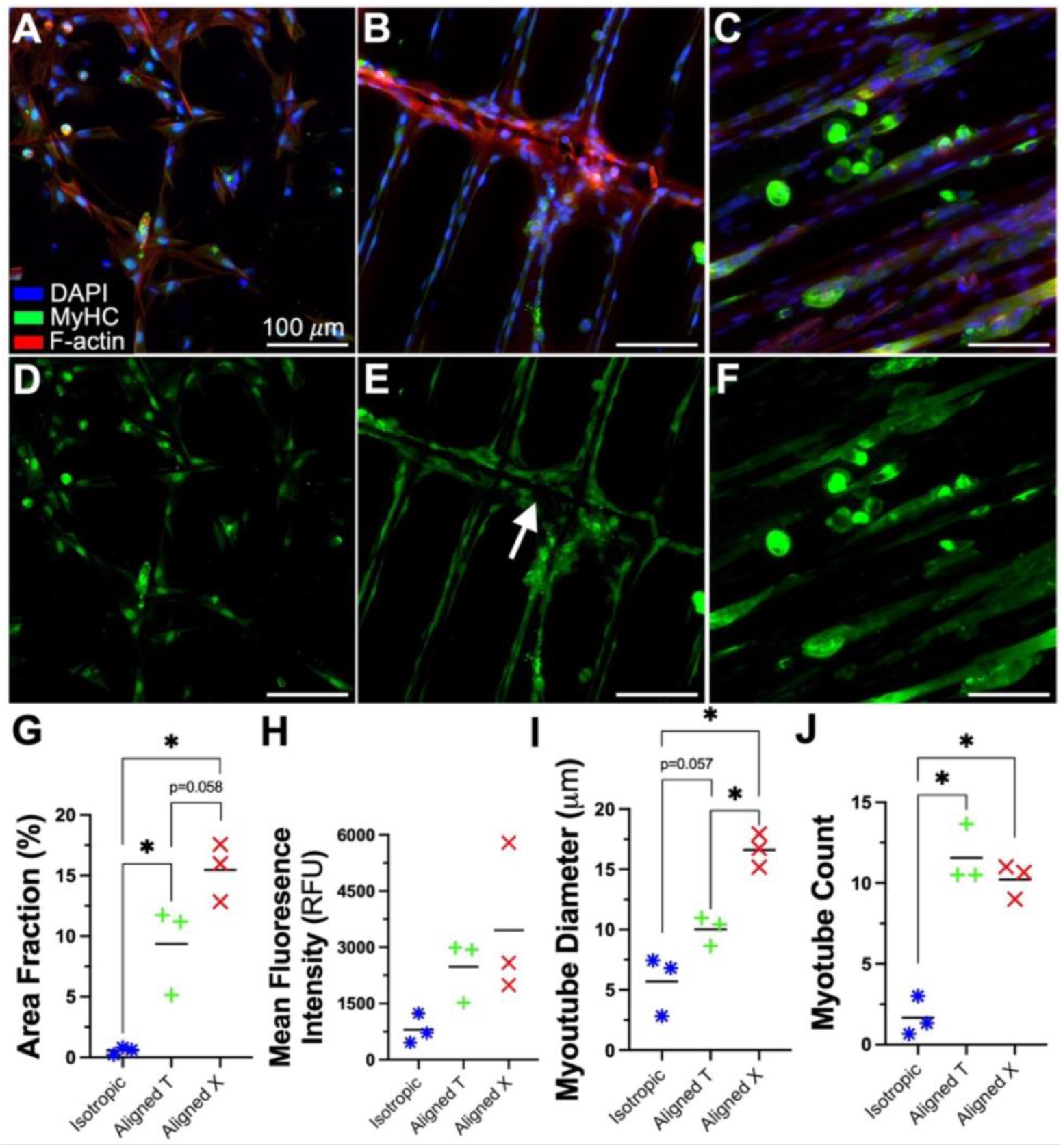
Composite constructs with Aligned X microfiber design had increased myotube formation. **A-C)** Immunofluorescence (IF) images of composite constructs with Isotropic (A), Aligned T (B), and Aligned X (C) microfiber scaffold design. The images show three colors with blue representing nuclei, green representing myosin heavy chain (MHC), and red representing F-actin. **D-F)** IF images of composite constructs in (A-C) only showing the green channel representing MHC. The acquisition settings included laser power of 3.0%, 2.4%, and 2.0% for the for the blue, green, and red excitation lasers, a gain of 849,578, 617.208, and 759.740 for the blue, green, and red emission channels, and a pixel dwell time of 2.576•10-7 seconds. **G)** Plot comparing the area fraction of myotubes across the three groups showed the Aligned X group had the highest myotube area fraction that was statistically different from the Isotropic constructs. **H)** Comparison of the mean fluorescence intensity of MHC across the groups. There were no statistical differences **I)** Analysis of myotube diameter across the three groups showed the Aligned X group had significantly larger myotube diameters than the Isotropic and Aligned T constructs. **J)** The number of myotubes in the Aligned T and Aligned X constructs was significantly higher than the Isotropic samples. The solid black line represents the mean of each distribution. Statistical significance was determined at p<0.05 with one asterisk. A one-way ANOVA was used with a Tukey’s post-hoc test.

## Discussion

This study introduced a novel skeletal muscle tissue engineering platform using microfiber-hydrogel composite constructs. The system uniquely integrates the advantages of high-precision additive manufacturing – through MEW – with hydrogel mediated cell seeding to enhance engineered tissues. The designed microfiber architecture effectively regulated the cellular organization from the microscale (∼10 µm) to the macro scale (∼1cm), modulated mechanical properties (Figure 2), and provided bulk structural stability (Figure 4). The hydrogel component improved cellular encapsulation efficiency, increasing cell retention, metabolic activity, and viability (Figures 5,6). Composite constructs also demonstrate design versatility through three distinct scaffold architectures that change cellular organization and significantly influence myogenic cell development (Figures 3,7).

Among these, the Aligned X scaffold architecture enabled constructs with improved myoblast differentiation and myotube maturation. Large scaffolds consisting of long fibrillar structures need structural reinforcements to maintain handling stability. Lack of reinforcements resulted in increased movement that lead to less cellular alignment (Supplementary Figure 1C,7). Perpendicular reinforcements have been previously utilized to address this problem^37,39,47^. Yet, the reinforcements have deleterious effects on cellular organization (Figure 3E,H and Supplementary Figure 5C,D), as they prevent long continuous myotube formation (Figure 7E). Perpendicular reinforcements were also found to influence hydrogel contraction that led to phase separation at the (Figure 4G). A recent approach with cardiomyocytes on MEW scaffolds reduced the amount of perpendicular reinforcements so the longitudinal direction of fiber walls remained continuous with highly sparse reinforcements^38^. This model utilized fibrin as the soft ECM component, and the cell-hydrogel constructs detached from the microfiber structures. By comparison, the Aligned X architecture permitted continuous alignment over large distances (Figure 4, Supplementary Figure 3) with minimal effect on cellular organization as cells conformed to the scaffold geometry (Supplementary Figure 5). Further, the Aligned X architectures significantly increased expression of MyHC and myotube diameters (Figure 7). This is partly due to the increased cellular continuity without barriers to long myotube formation. The increased myotube maturation could also be due to increased microenvironmental structural integrity, as the Aligned X microfiber scaffolds maintained their geometry more than Aligned T scaffolds (Supplementary Figure 2).

Scaffold mechanical properties were markedly tailored by architectural changes alone, resulting in more than two-fold from 3.89 ±0.575 MPa in the Isotropic group to 10.1 ±1.62 MPa in the Aligned X group (Figure 3B) – without changing scaffold mass. The magnitude of these moduli is similar to *ex vivo* tensile modulus of human levator ani muscles reported to be k = 4.69 ±2.91 MPa by Nagle *et al*.^48^, a linear scaling parameter of a power law constitutive model. Meanwhile, longitudinal extension of the extensor digitorum longus of rabbits, a hindlimb muscle, had a linear modulus of 447 ±97.7 KPa^49^. The mechanical properties of scaffolds can further be tailored by altering the microfiber diameter, the number of fibers, their spacing, or the polymer type. These scaffold variables therefore provide a large design space through which the mechanical properties can be adjusted to recapitulate specific tissue needs.

This study had several limitations. Myoblast seeding densities were relatively low compared to previous muscle engineering studies^38,50,51,55^. This could enable scalability with primary or human muscle progenitor cells that proliferate more slowly in future studies. Nonetheless, the current approach yielded constructs with cells populating most of the area along the microfiber walls by eight days of culture (Figure 3) and resulted in robust cellular differentiation by day 16 (Figure 7). Seeding of myoblasts on scaffold-only constructs resulted in clumping and inconsistent dispersion of cells. Prior similar approaches have attempted to perform chemical surface modifications to promote hydrophilicity^56^. By comparison, this study functionalized scaffold by overnight incubation with culture media that contains serum proteins, likely adsorbing onto the PCL microfibers. This may have influenced the viability and cellular retention of the scaffold-only group. The Isotropic, Aligned T, and Aligned X scaffold designs displayed different amounts of relative porosity in the axial direction (Figure 1), and the viability in the scaffold-only group seemed to correlate with the relative scaffold density (Figure 6B). The exclusion criteria of 30 nuclei per image resulted in a total of 3 samples discarded from the scaffold-only group while none were excluded from the composite samples (Figure 6). This suggests the data overestimate the cell count and viability of the scaffold-only group. Lastly, the aligned scaffold designs were not directly compared to other anisotropic scaffolds previously used in engineered muscle constructs. Thus, the high degree of alignment is only compared with the Isotropic scaffolds and control groups. Further studies are needed to delineate degrees of anisotropy and cellular alignment amongst state-of-the-art muscle constructs and examine their relevance to anatomic muscle fiber alignment.

The utilization of microfiber scaffolds as a fiber reinforcement to soft matrices advances the design space and capabilities for engineered tissue constructs with micron-scale precision. Using solid structures to prevent soft matrix contraction, and guide contraction along a particular axis have been utilized extensively in prior studies^50-54^. Likewise, microfiber scaffolds shown in this study dramatically altered cellular organization (Figure 3, Supplementary Figure 5), while stabilizing the bulk tissue geometry (Figure 4). MEW technology enables fabrication of micron-scale reinforcements over constructs larger than a centimeter. This could enable design of muscle fiber geometries with multipennate geometry, or curvature to recapitulate muscles that traverse a joint. Alternatively, it may be used to introduce spatial heterogeneities by manufacturing scaffolds with subregions that are not supported with microfibers to model structural deficits in muscle tissue or introduce a higher concentration of cross-bridge reinforcements in one location to model the musculotendon junction. These innovations allow for the tailorability of scaffold design across the cellular scale and bulk tissue scales and could advance engineered tissues by enabling greater design specificity.

## Acknowledgments

The authors acknowledge the following contributions: Dr. Genevieve Romanowicz for the development of seeding protocol and organization of the manuscript; Phillip Hernandez for technical help conducting the alamarBlue and picoGreen assays; Izzy Harker for implementing image processing scripts with python and cellpose; Kelly O’Neill for sharing seeding protocols and fabrication of electrospun membranes; Patrick Hall for laser cutting of electrospun membranes and aide in imaging gel contractions; Adam Fries of the Genomics and Cell characterization Core Facility at the University of Oregon for training on a Zeiss LSM confocal microscope; Kurt Langworthy from the Center for Advanced Materials Characterization in Oregon for assistance in perform scanning electron microscopy; the Knight Campus X-ray Imaging Facility and its Director, Angela Lin, for providing scan and image processing expertise and guidance. Scans performed with the Zeiss Xradia 620 Versa were made possible with equipment acquisition funding support from the M.J. Murdock Charitable Trust (Grant # SR-201812008). The research was funded by the Wu Tsai Human Performance Alliance and the Joe and Clara Tsai Foundation.

## Notes

Data availability statement: Data is available within the article and its supporting material, and also available upon request from the authors.

Conflict of Interest: A.R, I.L, P.D.D, R.E.G, and N.J.W have a provisional patent on the aligned X scaffold geometry manufactured using EHD printing. P.D.D and I.L. have an outside activity agreement with VivoTex LLC in which they serve as cofounders. The University of Oregon remains P.D.D’s primary place of employment and this research is attributed to the University of Oregon.

### Competing Interest Statement

A.R, I.L, P.D.D, R.E.G, and N.J.W have a provisional patent on the aligned X scaffold geometry manufactured using EHD printing. P.D.D and I.L. have an outside activity agreement with VivoTex LLC in which they serve as cofounders. The University of Oregon remains primary place of employment of Dr. Dalton and this research is attributed to the University of Oregon.

