## Supplementary Figures for "Melt Electrowritten Microfiber-Hydrogel Composite Scaffolds for Aligned Muscle Tissue Engineering"

Supplementary materials

### Supplementary Figure 1

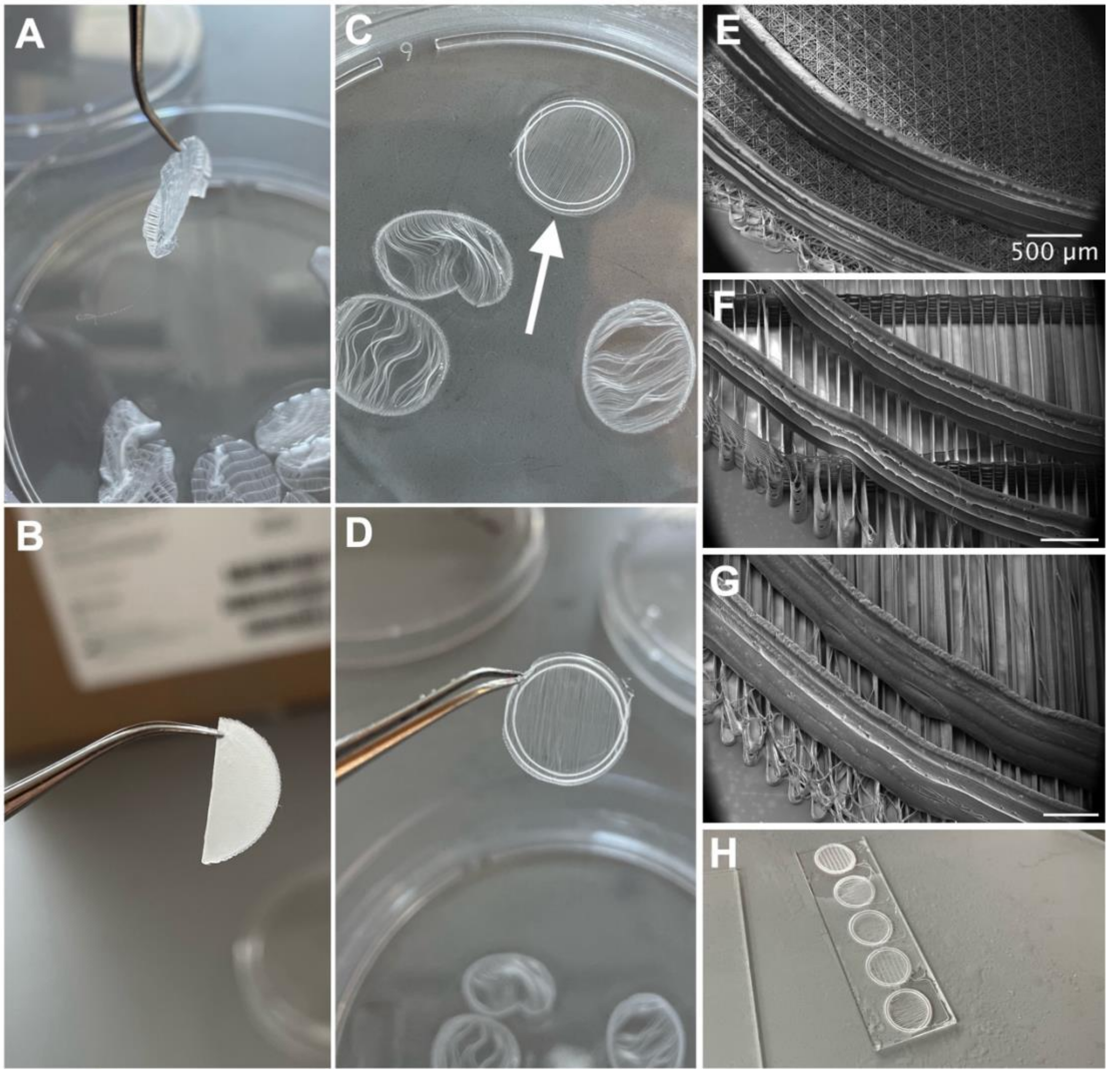


**Supplementary Figure 1.** Two Large, 70μm diameter, fibers reinforcing the scaffold along the perimeter improved its handleability. **A)** Aligned T scaffold without reinforcing rings wetted with ethanol and held with forceps. The scaffold contracts and folds to minimize the surface tension of the fluid. **B)** Isotropic scaffold without reinforcing rings wetted with ethanol and held with forceps folded in half. **C)** Aligned X scaffold with reinforcing rings (white arrow) placed next to aligned fibers without cross-bridge reinforcements or reinforcing rings. All scaffolds are wetted with ethanol, yet the non-reinforced scaffolds show considerable deformation, whereas the reinforced scaffold maintains its shape. **D)** Aligned X scaffold with reinforcing rings wetted with ethanol and handled with forceps maintained its shape. **E-G)** Scanning electron microscopy images of the large reinforcing rings on the Isotropic (E), Aligned T (F), and Aligned X (G) microfiber architecture designs. **H)** microfiber scaffolds with reinforcing rings printed on glass slides.

### Supplementary Figure 2

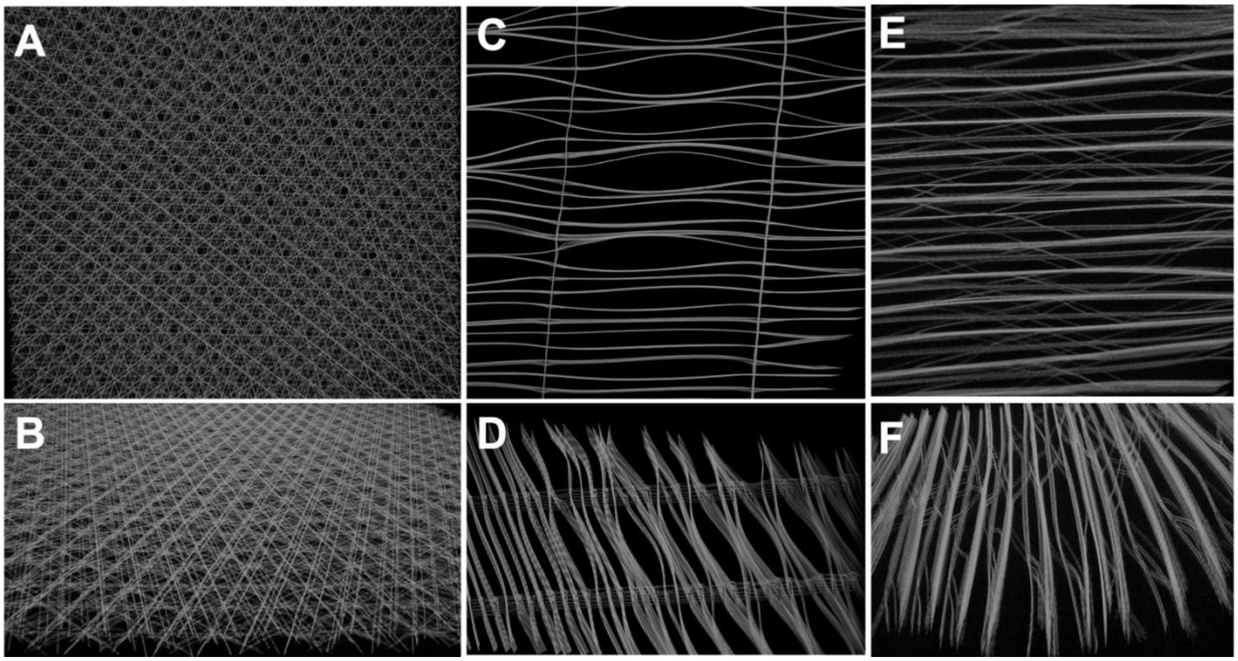


**Supplementary Figure 2.** Micro-computed tomography (µCT) images of microfiber scaffold architectures demonstrate the cross-bridge reinforcement improved the structural stability of aligned fibers. All scaffolds were imaged after wetting and allowing for a minimum of 3 hours of drying. **A-B)** Isotropic scaffold visualized via a top-view (A) and isometric off-angled view (B). **C-D)** Aligned T scaffold visualized in top and off-angled views. The aligned fiber tracks noticeably deform and portray a wavy pattern away from the perpendicular reinforcements. **E-F)** Aligned X scaffold visualized in top and off-angled views. The aligned fiber tracks appear to maintain their form better than the aligned T design.

**Supplementary Figure 3**

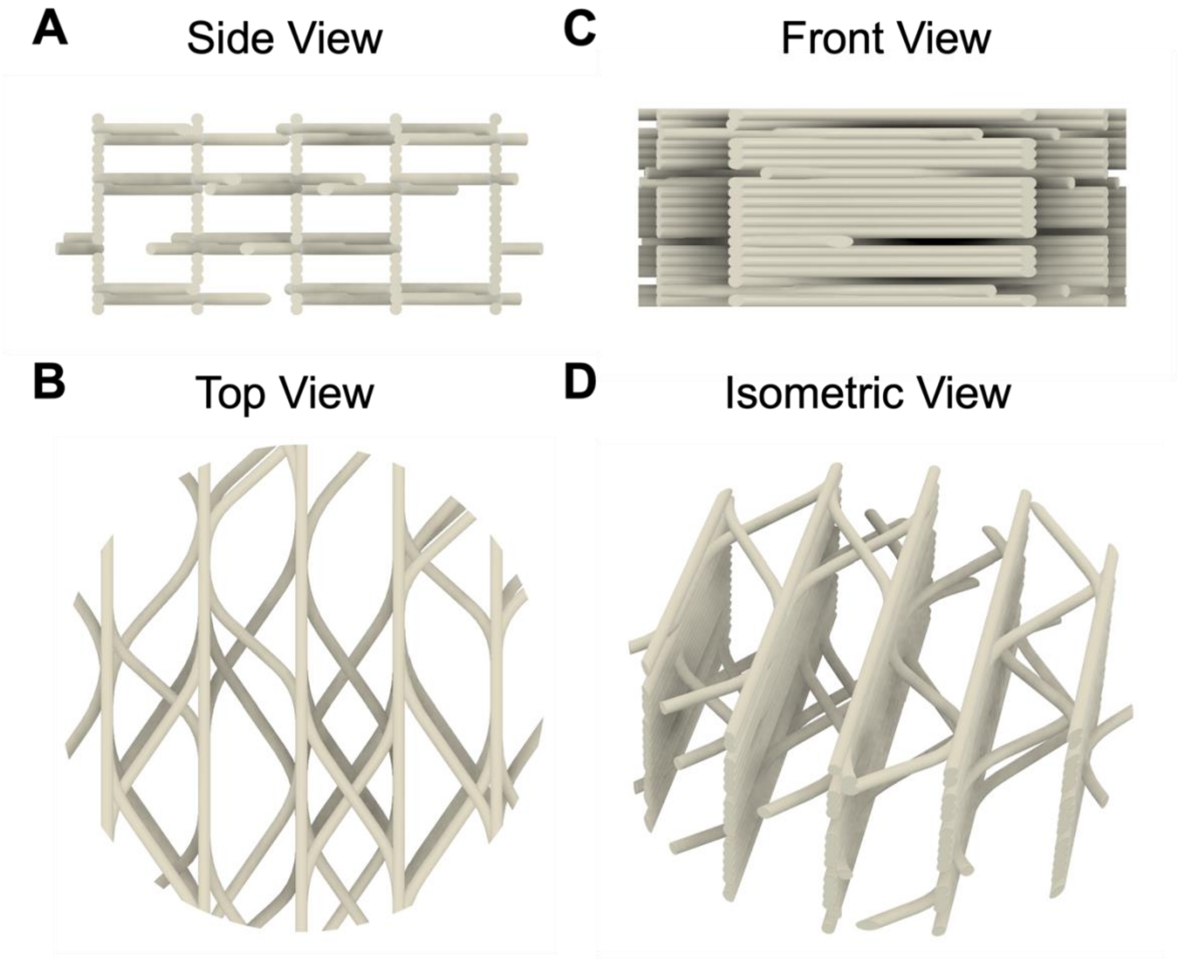


**Supplementary Figure 3.** Model representation of aligned X fiber architecture.

**Supplementary Figure 4**

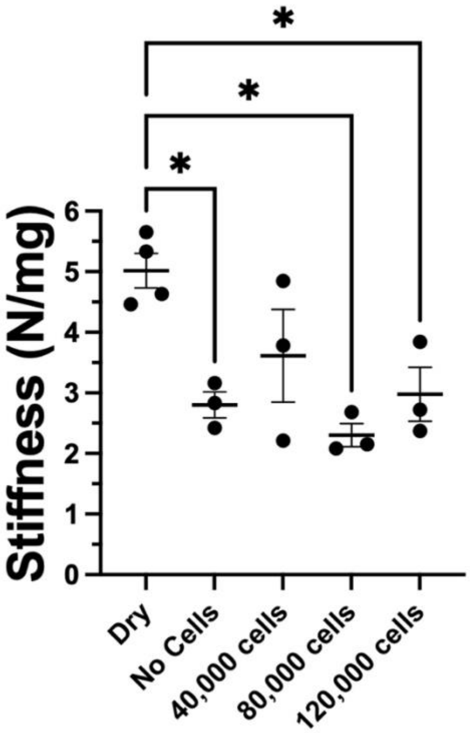


**Supplementary Figure 4.** Culturing cells on scaffolds did not influence their bulk-mechanical properties. Scaffolds of aligned X design were cultured with C2C12 myoblasts for 11 days with 40,000, 80,000, or 120,000 cells. Control groups included dry scaffolds that were never wetted or incubated, and acellular scaffolds incubated alongside the cellular scaffolds submerged in media. The groups did not include collagen hydrogel. After 11 days in culture, the constructs were allowed to dry for 1 hour and then tested under uniaxial tension as described in the methods section. Prior to testing, each construct's mass was recorded and used for normalization in place of width and thickness. The slope of the force-length curve was recorded and normalized to the mass of each sample. The results showed that constructs softened with incubation in media, but the presence or number of cells did not significantly influence the stiffness.

### Supplementary Figure 5

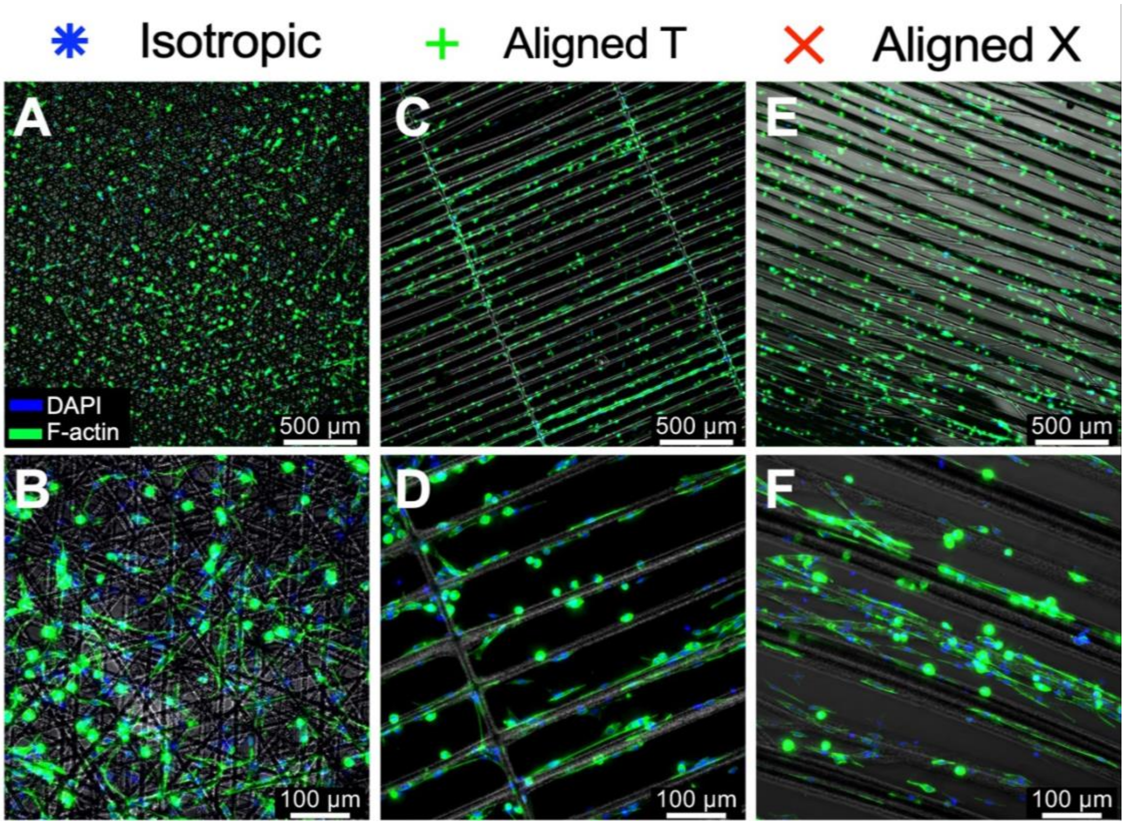


**Supplementary Figure 5.** C2C12 myoblasts adhered to microfiber scaffolds of composite scaffold-hydrogel constructs. All images are of composite constructs cultured to day 8. Immunofluorescence images showing the cellular cytoskeleton in green and nuclei in blue, overlayed on phase contrast images showing the microfiber scaffold in grayscale. **A-B)** Isotropic composite constructs acquired with a 5X objective (A) and a 20X objective (B). **C-D)** Aligned T Constructs showing 5X (C) and 20X images (D). **E-F)** Aligned X constructs in 5X (E) and 20X (F) magnification.

### Supplementary Figure 6

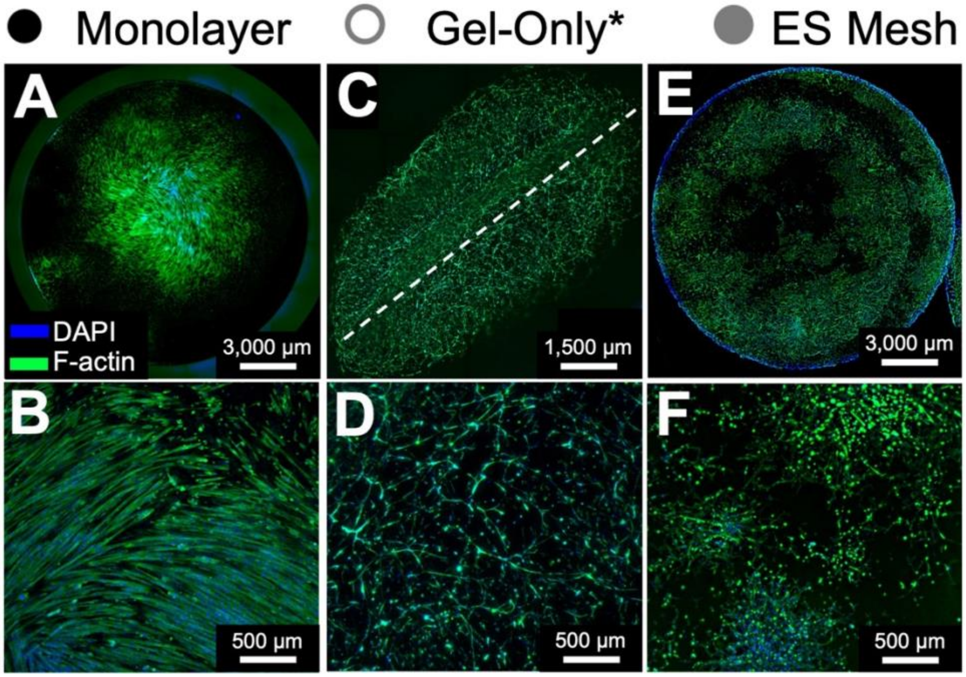


**Supplementary Figure 6.** C2C12 myoblasts cultured in control groups for comparison with composite microfiber scaffold-hydrogel composite constructs. The groups consisted of C2C12 myoblasts cultured in monolayer (A-B), hydrogel-only (C-D), and electrospun mesh (E-F).

### Supplementary Figure 7

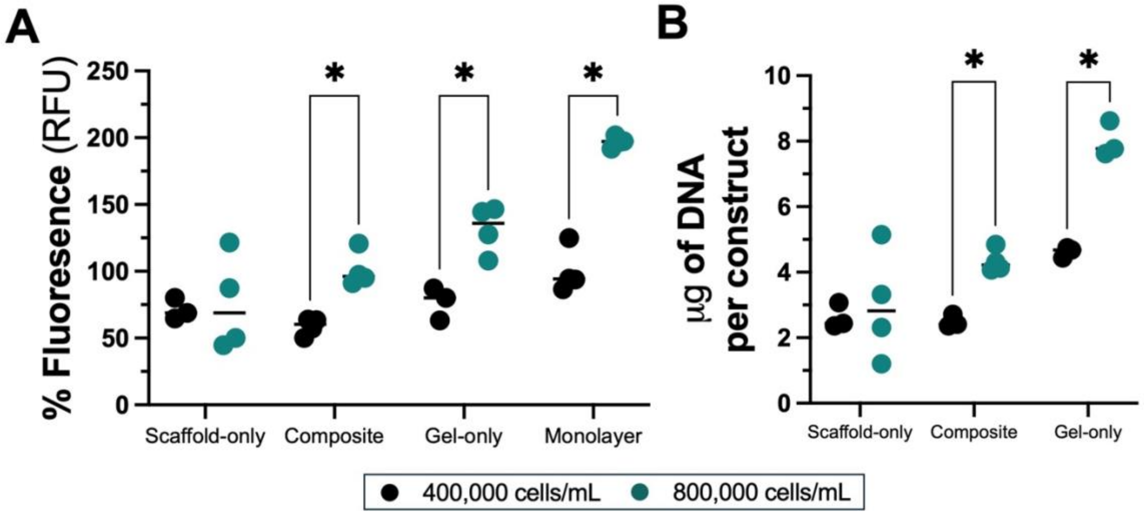


**Supplementary Figure 7.** Incorporation of a hydrogel enhanced metabolic activity and improved cellular seeding efficiency in composite constructs. This figure includes the information in Figure 5 with addition of control groups: Hydrogel-only (Gel-only) group and cell monolayer in alamarBlue (A), and hydrogel-only in picoGreen (B). **A)** Graph showing the results of alamarBlue assay comparing scaffold-only constructs and composite constructs 24 hours post seeding. The groups had two different effective seeding densities of 400,000 and 800,000 cells/mL. The composite constructs had increased signal from the lower to the higher seeding density, while the scaffold-only groups were not different across the seeding densities. **B)** Graph showing the results of picoGreen assay of the same groups in (A). Composite constructs had increased DNA mass with increased cell density, whereas the scaffold-only groups were not different from each other.

### Supplementary Figure 8

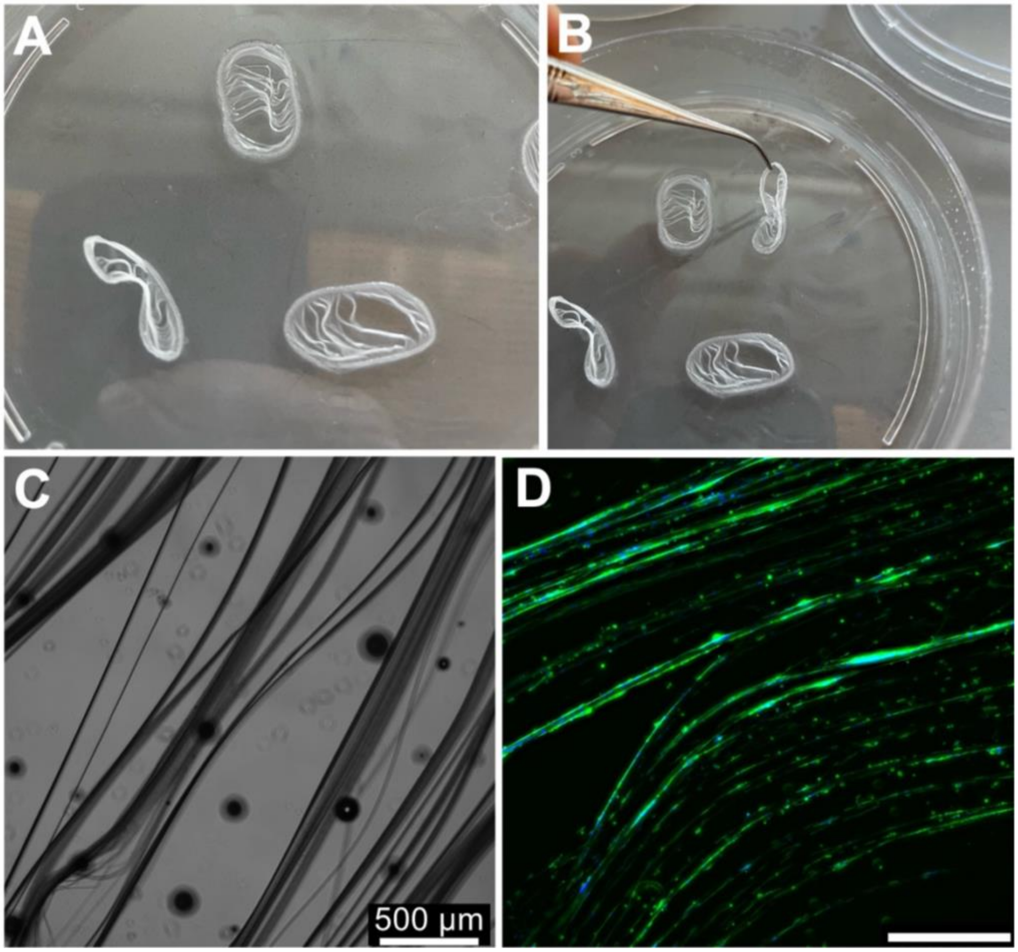


**Supplementary Figure 8.** Aligned microfibers scaffolds without reinforcements had considerable deformations. **A-B)** Aligned microfibers wetted with ethanol in a petri dish exhibit significant shape changes (A) and further deform when handling with forceps (B). **C-D)** Composite constructs utilizing aligned microfibers without reinforcements demonstrated in 20X images showed considerable dimensional instability via phase contrast (C), and immunofluorescence (D). Scale bar of 500 microns.
